# An epigenetic mechanism of azole tolerance facilitates acquired antifungal resistance in *Aspergillus fumigatus*

**DOI:** 10.64898/2026.03.16.712083

**Authors:** Sandeep Vellanki, Nathan DeMichaelis, Catherine Liao, Jason E. Stajich, Robert A. Cramer

**Author notes:** **Corresponding Author:** Robert A. Cramer, Ph.D.

## Abstract

Antibiotic tolerance paves the way for acquired resistance in bacterial pathogens. However, the mechanisms of tolerance and its evolutionary role in acquired resistance in pathogenic fungi, particularly molds, remains elusive. Here, we identified an <u>In</u>hibitor of <u>G</u>rowth domain protein (IngB) as a novel epigenetic regulator of azole tolerance in *Aspergillus fumigatus*. The loss of *ingB* promotes supra-MIC growth on agar surface despite susceptible MICs in standardized assays. Moreover, established Δ*ingB* biofilms are less susceptible to azoles *in vitro* and *in vivo*. Subsequent exposure of the tolerant strain to high azole concentrations resulted in rapid acquired resistance, most notably a frameshift mutation in a putative 20S proteasome maturation protein, UmpA, while the susceptible wildtype strain failed to acquire adaptive mutations. The data suggest that IngB-mediated tolerance provides an epistatic background for the emergence of azole resistance. Our work shows drug tolerance facilitates resistance emergence in a critical fungal pathogen.

**Importance:** While antimicrobial drug resistance causes a significant adverse effect on human health, drug tolerance can also lead to insufficient pathogen clearance, resulting in infection relapse. However, the mechanisms of antifungal drug tolerance and its evolutionary role in acquired drug resistance in pathogenic fungi, particularly the molds, remains elusive. We identified IngB as a novel regulator of azole tolerance in *Aspergillus fumigatus*. Importantly, loss of IngB leads to rapid azole drug resistance under azole-selective pressure. Our work identifies a novel regulator of antifungal tolerance and suggests antifungal drug tolerance can pave the way for resistance emergence in a critical fungal pathogen.

## Introduction

Drug resistance is a primary global reason for clinical therapeutic failures, leading to prolonged illness, deaths, and higher hospitalization costs (1). Often discussed in the context of cancer or bacterial infections, the continually emerging resistance to pathogenic fungi also poses a significant threat to humankind (2). One such emerging multidrug-resistant fungus, *Aspergillus fumigatus,* is a critical fungal pathogen and the causative agent of invasive and chronic aspergillosis (3, 4). Azoles, which target the Cyp51 enzyme in the ergosterol biosynthesis pathway, are the current frontline defence against filamentous fungal infections (5). However, their efficacy in vivo is compromised by acquired resistance driven by Cyp51 mutations and non-Cyp51 mechanisms (6). Besides resistance, clinical treatment failures also occurs with susceptible isolates by standard laboratory tests, (7) resulting in persistent infections (8). This suggests that alternative microbial mechanisms enable pathogenic fungi to overcome drug pressure.

Antifungal tolerance is defined as the ability of a microbial population to survive or grow at drug concentrations higher than the MIC, albeit slowly, without a change in MIC (9, 10). Tolerance mechanisms are far less well understood than resistance mechanisms in human pathogenic fungi with seminal studies largely focused on the pathogenic yeast (11–14). In bacteria, tolerance premediates resistance by persisting under antibiotic pressure long enough to acquire and fix stable mutations (15–18). It is unclear if tolerance is a prerequisite for azole resistance in *A. fumigatus* and if so, how a strain can transition from tolerance to formal resistance. Moreover, regulators of antifungal tolerance in pathogenic molds are ill-defined.

Given the critical role chromatin plays in sensing environmental changes and facilitating adaptation through transcriptional rewiring in response to stress (19, 20), we decided to investigate unexplored families of chromatin regulatory proteins in *Aspergillus fumigatus* for their role in azole drug responses. One such family is the <u>in</u>hibitory <u>g</u>rowth domain (ING; Pfam domain: PF12998) containing proteins that are conserved in all eukaryotes (21). *A. fumigatus* also possesses three proteins with an ING domain. Interestingly, one particular family member, AFU3G11940 (*ingB*), was mutated in a set of longitudinal isolates collected from a patient with cystic fibrosis (22). Moreover, in a series of longitudinal collected isolates from a patient with Chronic Granulomatous Disease (CGD) with reduced susceptibility to azoles, *ingB* variant alleles were also detected (23). Here, we found that loss of *ingB* results in tolerance to azoles *in vitro* and *in vivo* in a murine model of invasive pulmonary aspergillosis. Surprisingly, we observed that this azole tolerant mutant acquires resistance when exposed to azoles. Notably, a frame shift mutation in the putative 20S proteasome gene (*umpA*) arises in *ingB* null mutants and results in reduced growth and conidiation, increase in vegetative mycelium, and resistance to all azoles, a characteristic typical of isolates collected from patients with chronic aspergillosis (6). Therefore, our study addresses a significant knowledge gap by identifying novel gene(s) and mechanisms related to *A. fumigatus* azole drug tolerance and its role in the emergence of drug resistance.

## Results

### Loss of *ingB* results in tolerance to triazoles

The genetics strategy employed for the generation of Δ*ingB* and Δ*ingB*^Rec^ are detailed in Methods and the validation of strains is shown in **Supplementary Fig. 1**. Loss of *ingB* resulted in a modest reduction in colony biofilm growth on glucose minimal medium and minimal morphological changes that were both reconstituted with re-introduction of the wild-type allele **(Fig. 1)**. We next tested Δ*ingB* susceptibility to azoles using the standardized CLSI microdilution method. Loss of *ingB* did not result in a change in susceptibility to voriconazole, itraconazole, or posaconazole (Table 1). However, surprisingly, while growth in the presence of 0.2 µg/ml voriconazole in agar resulted in about a 50% reduction in colony diameter of the WT and reconstituted strains, and no growth was observed at 0.4 µg/ml or 0.8 µg/ml, which are concentrations above the MIC, the Δ*ingB* strain only showed a ∼20% reduction in colony diameter in the presence of 0.2 µg/ml voriconazole, a 50% reduction at 0.4 µg/ml, and a 90% reduction at 0.8 µg/ml voriconazole **(Fig. 1)**. These data suggest that *ingB* is an important mediator of azole tolerance.

**Figure 1.**
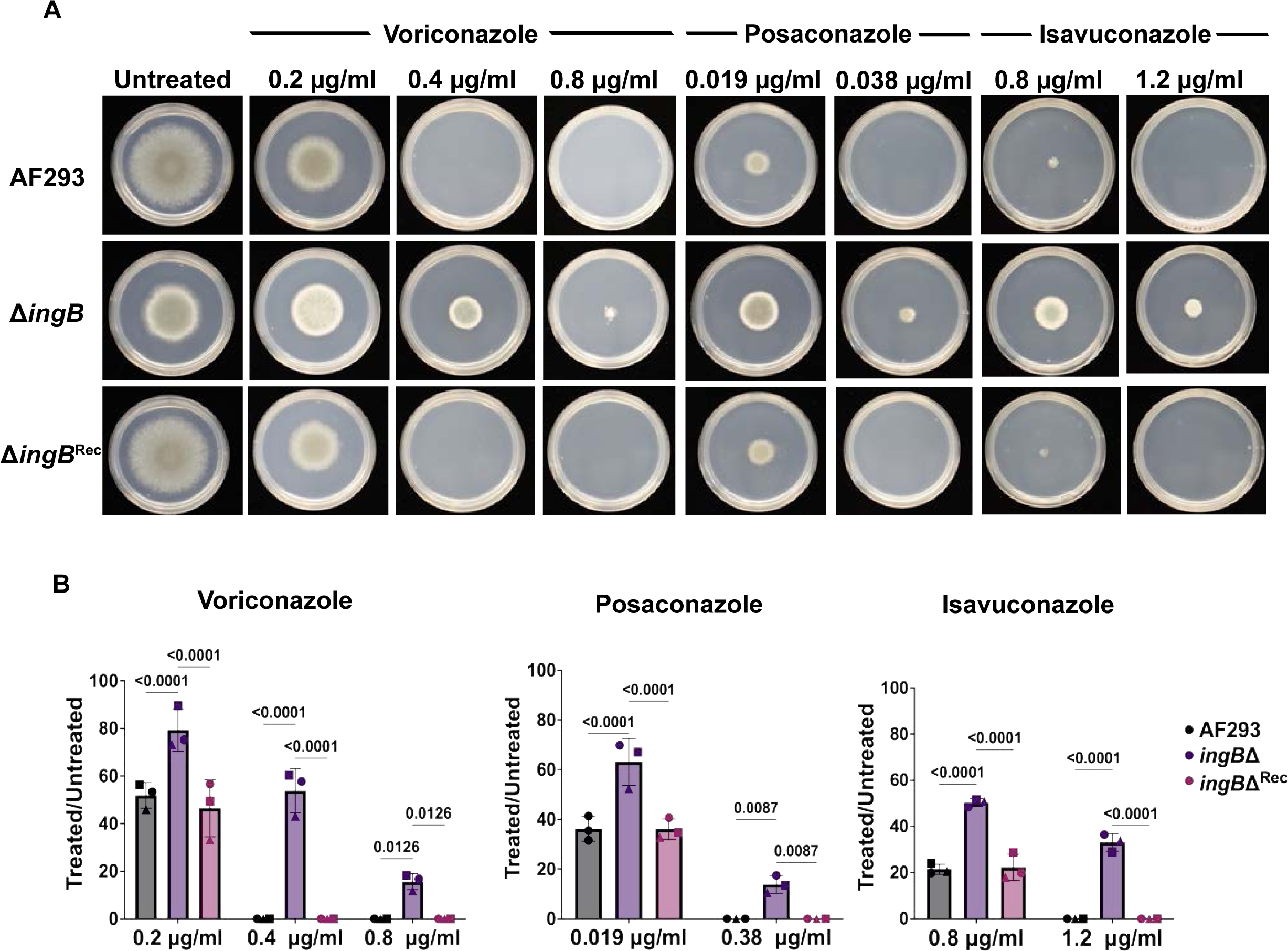
Loss of *ingB* results in tolerance to azoles on solid agar plates. A) 1000 conidia were inoculated in GMM alone or in the presence of voriconazole, posaconazole or isavuconazole. After 72 hrs, the radial growth was measured, and the images were taken. B) Two Way ANOVA for row factor was significant for voriconazole, posaconazole, and isavuconazole (P<0.0001). Dunnet’s multiple comparison test was used to compare each group with the mutant. The p-value is indicated on the graph. Data is representative of 3 biological replicates (indicated by symbols).

**Table 1.** Minimum Inhibitory Concentrations of strains generated in the study.

| Strains | Voriconazole | Itraconazole | Posaconazole | Isavuconazole | Amphotericin B |
| --- | --- | --- | --- | --- | --- |
| AF293 | 0.25 | 0.25 | 0.031 | 1 | 1 |
| *ΔingB* | 0.5 | 0.5 | 0.031 | 2 | 1 |
| *ΔingB*<sup>Rec</sup> | 0.25 | 0.25 | 0.031 | 1 | 1 |
| *ΔumpA* | 2 | >8 | 0.125 | 4 | 2 |
| *ΔumpA*<sup>Rec</sup> | 0.25 | 0.25 | 0.031 | 1 | 1 |
| *ΔumpA*<sup>Ser55fs</sup> | 2-4 | >8 | 0.125 | 4 | 2 |
| *ΔingB*<br>*umpA*<sup>Ser55fs</sup><br>(colony 11) | 4 | >8 | 0.125 | 8 | 2 |
| colony 11<br>*umpA*<sup>AF293</sup> | 0.5 | 0.5 | 0.031 | 2 | 1 |

To test whether this tolerance phenotype was specific to voriconazole, we conducted the same assay with other clinically relevant azoles. In the presence of 0.019 µg/ml posaconazole, the WT and reconstituted strains showed approximately a 75% reduction in growth. In contrast, the Δ*ingB* strain showed a 40% reduction in growth **(Fig. 1)**. At the concentration above the MIC, the WT and reconstituted strains showed no growth while the Δ*ingB* had persistent growth. The MIC for isavuconazole is 1 µg/ml (**Table 1**). Consistent with observations with the above data, at a sub-MIC concentration of 0.8 µg/ml isavuconazole, the WT and reconstituted strains showed approximately an 80% reduction in growth, whereas Δ*ingB* showed a 50% reduction in growth **(Fig. 1)**. At concentrations slightly above the MIC (1.2 µg/ml), 70% growth reduction was observed in Δ*ingB*, and no growth was observed in WT and the reconstituted strains. Similar observations were made with itraconazole **(Supplementary Fig. 2)**. Taken together, our data suggest that loss of *ingB* not only results in a strain with less susceptibility to azoles than WT at sub-MIC concentrations but one that can also grow at azole concentrations above the MIC on solid medium, despite no change in MIC. We conclude that loss of *ingB* leads to azole tolerance.

### Δ*ingB* biofilms are less susceptible to azoles *in vitro* and *in vivo*

Previous work from our group and others has shown that the biofilm growth stage significantly reduces antifungal susceptibility (24, 25). We wondered if a tolerant strain would also possess an alternation in biofilm drug susceptibility. We first compared the biofilms of Δ*ingB* with those of WT to ensure that biofilm formation and growth in this model are similar. As presented in **Fig. 2A**, Δ*ingB* biofilm morphology is similar to the WT (AF293) and the reconstituted strain at 16 hrs of development. The biofilm morphology is characterized by adherence to the base of the substrate, and hyphal growth along the XY and Z-axis over time. Additionally, no significant differences in biofilm total biomass were observed between WT, Δ*ingB*, and the reconstituted strain **(Fig. 2B)**. Therefore, we next tested biofilm antifungal susceptibility in the presence and absence of *ingB*.

**Figure 2.**
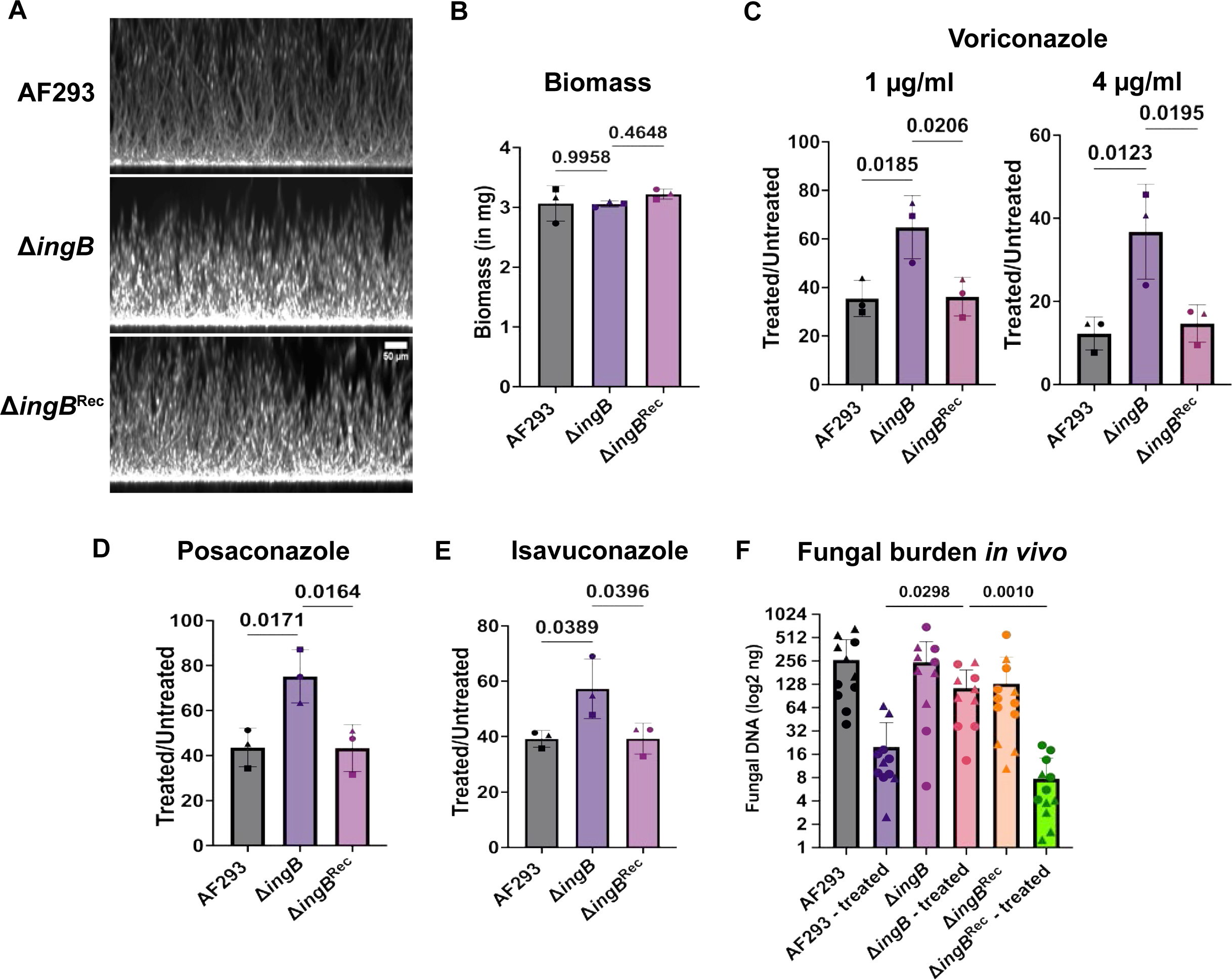
Loss of *ingB* results in reduced biofilm susceptibility to azoles *in vitro* and *in vivo*. A. Representative images of 16-hr biofilms stained with calcofluor white. Scale bar is indicated on the graph. B) No difference in biomass was observed among the groups at 16 hours, before drug treatment. For each technical replicate, biomass from two wells of a 6-well plate was pooled. One way ANOVA was not significant (P=0.4912). The experiment was repeated three times independently (indicated by symbols) with three technical replicates in each experiment. Dunnett’s multiple comparison test was used to compare each group with the mutant, and the *p*-value is indicated in the graph. For drug susceptibility assays, 16-hour biofilms were treated with C) 1 or 4 µg/mL voriconazole, D) 0.2 µg/mL posaconazole, and E) 2 µg/mL isavuconazole for 4.5 hours. The medium was then removed and replaced with fresh minimal growth medium. After 16 hours, biomass was collected. Treated-to-untreated biomass ratios are shown in the graphs for each concentration. The experiment was repeated three times independently (indicated by symbols). One-way ANOVA was significant for voriconazole at 1 µg/mL (*P* = 0.0166) and 4 µg/mL (*P* = 0.0128), and for posaconazole (*P* = 0.0141). Dunnett’s multiple comparison test was used to compare each group with the mutant, and the *p*-value is indicated in the graph. E) Loss of *ingB* results in reduced susceptibility to voriconazole *in vivo.* Four-to-six-week-old CD-1 mice were immunosuppressed, infected, and treated with voriconazole. Fungal burden was assessed through quantitative PCR (qPCR) quantitation of *A. fumigatus* 18S rDNA. The experiment was repeated two times, as indicated by symbols. The Kruskal-Wallis Test was significant (*P* < 0.0001). Dunn’s multiple comparison test was used to compare treated groups, and the *p*-value is indicated on the graph.

As presented in **Fig. 2C**, treatment with 1 µg/ml voriconazole resulted in a ∼65% reduction in biofilm growth at the tested time point in the WT and the reconstituted strain, whereas a significant reduction in susceptibility was observed in Δ*ingB* biofilms (∼40% decrease in biomass growth, compared to untreated). At the large 4 µg/ml dose of voriconazole, compared to untreated controls, the WT and the reconstituted strain had a ∼90% growth reduction while Δ*ingB* had a ∼65% reduction. Similar results were observed with posaconazole (**Fig. 2D**). The WT and the reconstituted groups had a ∼60% reduction in biomass growth when compared to untreated controls. However, Δ*ingB* had only a 30% reduction. The Δ*ingB* biofilms are also less susceptible to isavuconazole (**Fig. 2E)**. Taken together, Δ*ingB* tolerance to azoles is not dependent on its morphological form or the growth stage, as both Δ*ingB* conidia and biofilms show reduced susceptibility to azoles. As biofilm growth is relevant *in vivo,* and we wondered if the *ingB* azole tolerance would be releveant in a pre-clinical animal model, we next tested whether loss of *ingB* impacted fungal burden in a murine model of invasive aspergillosis treated with voriconazole.

The fungal burden between the groups not treated with voriconazole across strains was similar, suggesting that *ingB* is dispensable for fungal fitness and disease progression in this murine model (**Fig. 2F**). Loss of *ingB* also did not alter survival outcomes in triamcinolone model of invasive aspergillosis (**Supplementary Fig. 3**). However, mice challenged with the WT and the reconstituted strains and treated with voriconazole resulted in a ∼16-fold decrease in fungal burden. In contrast, mice challenged with Δ*ingB* and treated with voriconazole only resulted in 2.5-fold decrease in fungal burden (**Fig. 2F**). Consistently, Δ*ingB*-treated groups show statistically significantly higher fungal burden than the WT or reconstituted group treated with voriconazole (**Fig. 2F**). Therefore, consistent with the observations made *in vitro,* loss of *ingB* favours survival and growth under voriconazole treatment *in vivo*.

### Loss of *ingB* leads to repression of genes involved in ergosterol biosynthesis and an increase in iron starvation response

We next explored the potential mechanism behind the IngB mediated azole tolerance phenotype. A previous study identified IngB as part of the Nua3 Histone Acetyltransferase complex, which catalyses the acetylation of H3K9 and H3K14 (26). IngB orthologs are also found to be part of the same complex in other fungal species (27–29). Given its role in gene regulation and expression, we utilized RNA-seq to understand the differences in gene expression between Δ*ingB* and WT. The loss of *ingB* resulted in a significant change in the transcript abundance of 1,225 genes (p < 0.05), approximately 12% of the annotated genes in the *A. fumigatus* genome. The predominant number of transcripts in the mutant are reduced compare to the wild-type suggesting that IngB likely functions primarily as a positive regulator of gene expression. The list of all gene CPM values and fold change expression values is detailed in **Supplementary Tables 1-3**.

Several key genes involved in *A. fumigatus* growth, metabolism, stress response, and secondary metabolism, whose transcripts were significantly altered in the absence of *ingB*, are highlighted in **Fig. 3A**. We focused on pathways and mechanisms known for a role in pathogenesis and drug susceptibility. Among the genes with increased transcript levels, were several members of the heat shock protein (HSP) family, which play crucial role in mitigating stress and enhancing antifungal resistance (30). Several transcripts of genes encoding putative Glutathione S-transferases, which are generally involved in cellular detoxification and protection against oxidative stress (31), were also upregulated in the Δ*ingB* mutant (**Fig. 3A**). We also observed a modest increase in transcripts of *mdr1* (32) and *atrF* (33) drug transporters (**Fig. 3A**).

**Figure 3.**
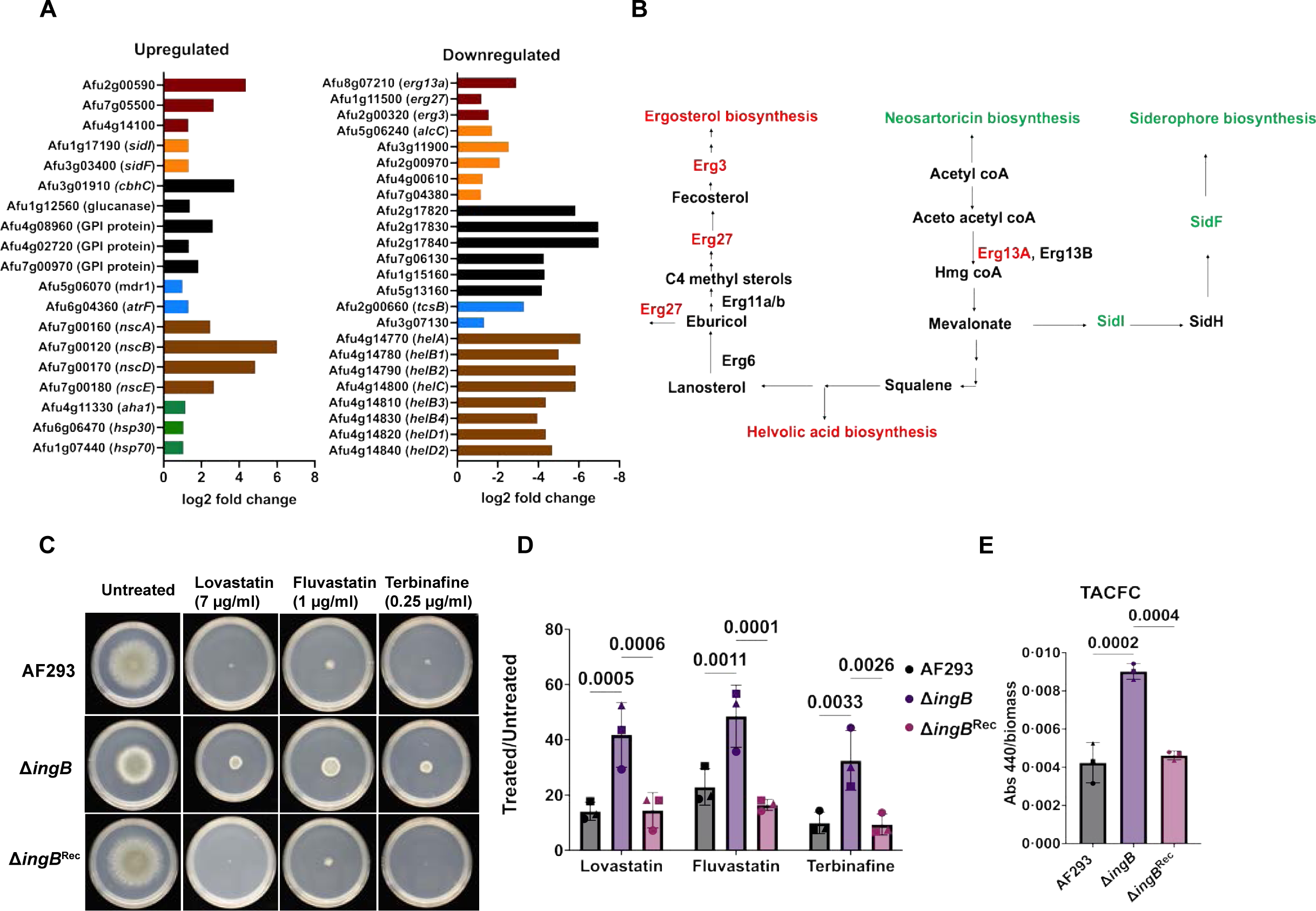
Deletion of *ingB* results in repression of genes in ergosterol biosynthesis and increased iron starvation response. A) Comparison of DEGs in Δ*ingB* vs WT. RNA-seq was performed on 16-hr biofilms of AF293 and Δ*ingB*. There were 1225 DEGs between Δ*ingB* and AF293. Of these, the expression of 536 genes was significantly changed by at least 2-fold, and their cpm values were more than 10. Select upregulated and downregulated genes (2-fold difference, p<0.05, cpm>10) in the mutant are shown in the figure: Upregulated – genes involved in putative glutathione S-transferases (dark red), siderophore biosynthesis (orange), cell wall (black), drug transporters (blue), neosartoricin biosynthesis (brown), and heat-shock protein genes (green). Downregulated – genes involved in ergosterol biosynthesis (dark red), alcohol dehydrogenases (orange), membrane integrity (black), two-component system (blue), and Helvolic acid biosynthesis (brown). B) The downregulation (indicated in red), of *erg13a*, *erg3*, and *erg27* alongside an increase (indicated in green) in siderophore biosynthesis genes and neosartoricin biosynthesis was observed. C) The conidia of each strain were inoculated on plates containing Lovastatin, Fluvastatin, Terbinafine. Radial growth was measured, and plate images were taken. D) Treated to untreated ratios are shown. Two Way ANOVA was significant (*P=0.0116). Dunnet’s multiple comparison test was used to compare each group with the mutant, and the p-value is indicated in the graph. The data shown is representative of 3 bioreps (indicated by symbols). E) Each strain was cultured in flasks containing iron-free GMM flasks. The flasks were incubated at 37℃, 5% CO2, and 200 rpm. TACFC production per weight of biomass was calculated. The data shown is representative of 3 bioreps (indicated by symbols). One way ANOVA was significant (P=0.0002). Dunnet’s multiple comparison test was used to compare mutant with WT and the recon groups. The p-value is indicated on the graphs.

Among the genes with decreased transcript levels in the absence of *ingB* are several putative genes encoding proteins involved in maintaining cell membrane integrity (**Fig. 3A**). Transcripts of two-component signal transduction systems, which perceive extracellular signals and activate downstream response pathways, were also significantly reduced in the *ingB* null mutant (34). Genes encoding transcripts related to primary and secondary metabolism were altered including those involved in the mevalonate pathway, several alcohol dehydrogenases, and the helvolic acid pathway (**Fig. 3B**).

As acetyl-CoA is a key building block for fungal sterols (35), we were particularly intrigued with transcripts related to acetyl-CoA altered with loss of *ingB.* Acetyl-CoA acts as a precursor to several primary and secondary metabolism pathways. As shown in Fig. 4B, acetyl-CoA acts as a precursor to the mevalonate pathway (35) and polyketide synthesis (36). Transcripts of the genes involved in neosartoricin biosynthesis were increased in the absence of *ingB*; transcripts of the early mevalonate pathway gene *erg13A* were significantly decreased. Mevalonate serves as a precursor to the ergosterol pathway, or it can be directed towards the synthesis of siderophores (37). Interestingly, transcripts of genes involved in siderophore biosynthesis, *sidI* and *sidF*, were significantly increased with *ingB* loss by 2.71-fold (**Fig. 3A**). With regard to helvolic acid synthesis, squalene, also a sterol precursor, is converted to (3S)-2,3-oxidosqualene that can be incorporated into the ergosterol biosynthesis pathway or utilized for helvolic acid biosynthesis (38). Perhaps correspondingly, transcripts of all genes involved in helvolic acid biosynthesis and transcripts of several genes in the ergosterol pathway were decreased (**Fig. 3B**). Together, the data suggest that IngB is a global regulator of primary and secondary metabolism, stress responses, and cell membrane homeostasis in *A. fumigatus*.

**Figure 4.**
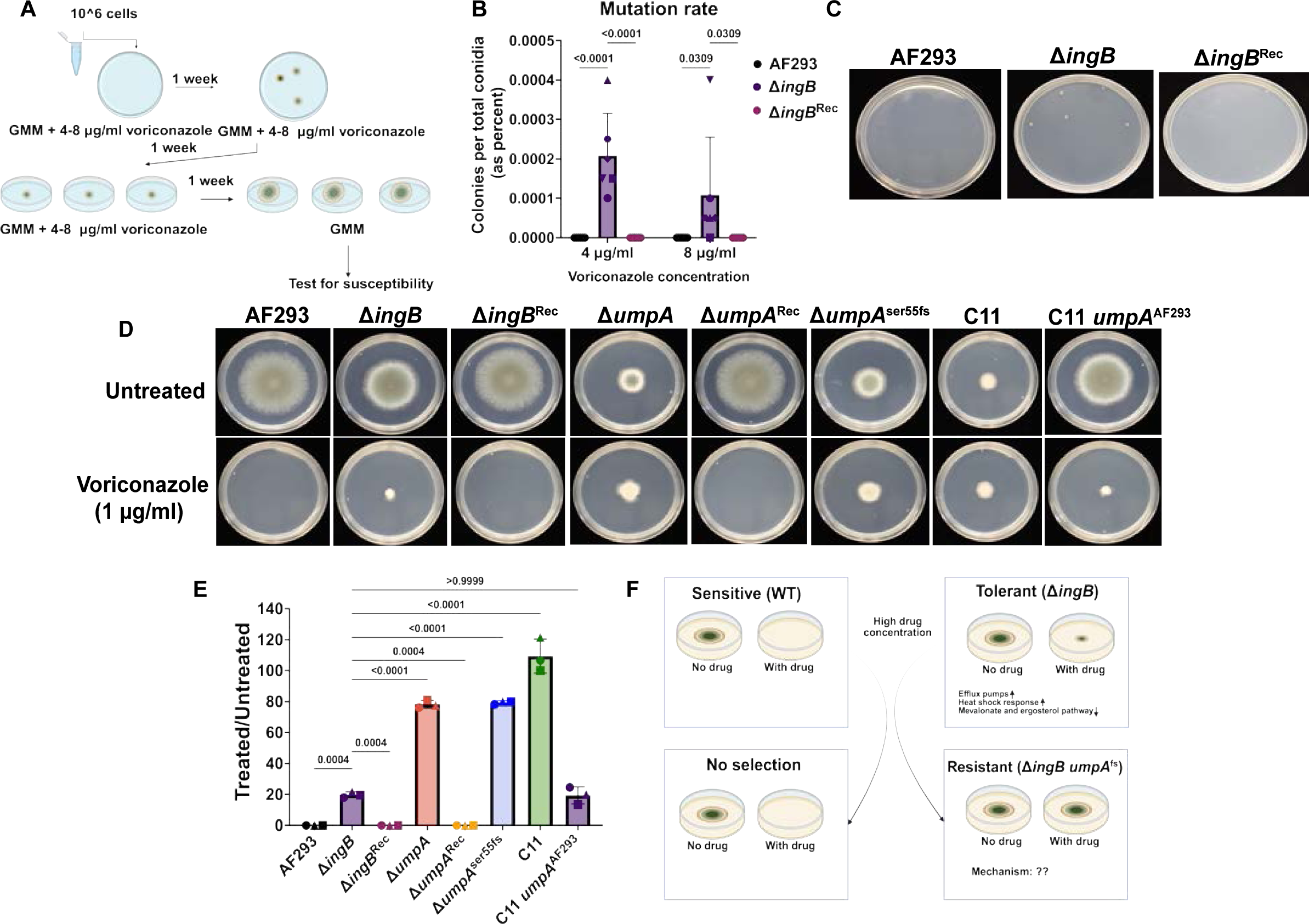
Loss of *ingB* results in selection of azole resistant strains with a frameshift mutation in *umpA*. A) The experiment design for selection of azole resistant strains. B) The mutation rate was calculated by total number of colonies per total population of conidia at two different voriconazole concentrations. Each point represents a different biorep. Two-way ANOVA for column factor (strain comparison) was significant (p<0.0001). For the row factor (different concentrations) no statistical significance was observed (p=0.1865). Dunnett’s multiple comparison test was used to compare groups at both the concentrations at the p-value is indicated on the graph. D) *umpA^Ser55fs^*drive azole resistance. The conidia of strain were spotted on GMM containing 0 or 1.5 µg/ml voriconazole for 72 hrs and radial growth was measured. E) Quantification of treated to untreated ratios of plates shown in D. One way ANOVA was significant (p<0.0001). Dunnett’s multiple comparison was used to compare each group with Δ*ingB*, and the p-value is indicated on the graph. Each biological replicate is indicated by a symbol in the graph. F) Azole tolerance accelerates the selection of acquired resistance. Drugs such as azoles inhibit the growth of a sensitive *A. fumigatus* strain (WT). However, a tolerant strain (Δ*ingB*) is less susceptible to azoles, despite no change in its MIC. This tolerance response is characterized by modest upregulation of stress responses such as heat shock proteins and efflux pumps, and downregulation of genes involved in the mevalonate and ergosterol pathways. When both the sensitive and the tolerant strains are exposed to a high concentration of azoles, this results in the killing of all sensitive cells. However, a sub-population of the tolerant population can gain mutations to become resistant, thereby providing a selection advantage. The model figure was created in Biorender. Cramer, R. (2026) https://BioRender.com/d1tbpn4.

### Alterations in mevalonate pathway transcripts likely drive Δ*ingB* tolerance

We hypothesized that loss of *ingB* results in global transcriptional rewiring leading to a constitutively active signal for iron starvation that forced a critical metabolic trade-off by shunting mevalonate pathway precursors away from ergosterol biosynthesis toward siderophore production. Consistent with this hypothesis, treating WT and the reconstituted strain with 7 µg/ml lovastatin, which targets HMG-CoA reductase, resulted in approximately 85-90% reduction in growth, whereas Δ*ingB* showed only a 60% growth decrease (**Fig. 3C and Fig. 3D**). Similar observations were also made with fluvastatin, which also targets HMG-CoA reductase, where Δ*ingB* is statistically significantly less susceptible than the WT and the reconstituted strain (**Fig. 3C and Fig. 3D**). Additionally, treatment with terbinafine resulted in about 90% reduction in colony size in the WT and the reconstituted strain, with a 70% reduction in colony size observed with Δ*ingB* (**Fig. 3C and Fig. 3D**). Importantly, susceptibility to Amphotericin B was not altered in Δ*ingB* (**Supplementary Fig. 4**). Taken together, these data suggest loss of *ingB* reduces flux through the ergosterol biosynthesis pathway but does not grossly change total ergosterol accessible to amphotericin B. Our hypothesis also predicts an increase in siderophore production. We therefore next compared TAFC siderophore production among the strains. As shown in Fig. 4E, Δ*ingB* produces significantly more TAFC per gram of biomass, compared to WT and the reconstituted strain. Taken together, these data support the transcriptomics analysis and indicate that loss of *ingB* likely perturbs the flux of key metabolites involved in production of the antifungal drug target, ergosterol.

### Δ*ingB* mediated tolerance drives selection of acquired drug resistance

As a putative loss of function *ingB* allele was discovered in a series of azole resistant isolates from a person with chronic granulomatous disease, and loss of *ingB* in our studies promotes azole tolerance likely through modulation of fungal metabolism associated with sterol production, we asked whether the tolerant *ingB* strain would be prone to development of true antifungal resistance. To address this significant knowledge gap, we conducted a selection experiment where we inoculated the parent AF293 (sensitive), Δ*ingB* (tolerant), and reconstituted strain (sensitive) at voriconazole concentrations above the MIC (**Fig. 4A**). As presented in **Fig. 4B and C**, despite a high inoculation concentration of conidia, we did not recover any colonies from the sensitive strains over 6 biological repeats, while dysmorphic but growing colonies were recovered from Δ*ingB* inoculated plates. The approximate mutation rate for *ingB* under this selection was calculated by the number colonies that emerged per total conidia inoculated in each biological replicate (39). A mutation rate of 0.002% and 0.001% was observed at 4 and 8 µg/ml respectively in Δ*ingB*. Colonies obtained on the Δ*ingB* drug plates were compact and appeared reddish. When these colonies were patched onto GMM with no drug, most colonies are compact and have increased vegetative mycelium (**Supplementary Fig. 5A**). All the colonies that were tested were resistant to voriconazole (**Supplementary Fig. 5B**). Therefore, our data suggest that tolerance conferred by the absence of IngB promotes the emergence of antifungal resistant mutants.

### *umpA^ser55fs^* is a novel mutation that confers triazole resistance

To determine how the tolerant strain (Δ*ingB*) became resistant, we utilized whole-genome sequencing on five independent colonies that exhibited resistance to azoles following exposure (**Fig. S5**). Colony 1 had a frameshift mutation in Afu5g08450 (UBA_TAP-C-like protein), colony 9 had a frameshift mutation in Afu5g06790 (NTF-2 domain protein), and colony 11 had a frame shift mutation in Afu5g10740, which is a conserved ortholog of the proteasome maturation factor *umpA* in fungi. We decided to investigate further if the *umpA^ser55fs^* is conferring azole resistance. We generated a Δ*umpA* in the WT background and then reconstituted the strain with the WT locus or with the colony 11 locus (which contains the *umpA^ser55fs^* allele) (see methods). Alternatively, we also reconstituted colony 11 with the WT *umpA* allele from AF293. Strain validation data are shown in **Supplementary Fig. 1**.

If the frameshift mutation results in a loss of function, we would expect the null mutant to exhibit the same phenotype as the evolved allele. Conversely, restoring the wild-type allele in colony 11 should restore azole susceptibility. Consistent with our hypothesis, both *umpA^ser55fs^* and the Δ*umpA* strains are resistant to voriconazole, itraconazole, posaconazole, and isavuconazole (**Table 1**). We did not observe a change in MIC with Amphotericin B. As expected, restoring the functional copy of *umpA* in colony 11 restored antifungal susceptibility to Δ*ingB* levels (**Table 1**).

We also adapted a second plate-based agar assay to test for azole susceptibility (**Fig. 4D and 4E**). As expected, we did not observe any growth of the WT and reconstituted strains on plates containing 1 µg/ml voriconazole. Interestingly, although colony 11, *umpA^ser55fs^,* and the Δ*umpA* strains have the same MIC, colony 11 exhibits more growth in the presence of high concentration of azoles. As expected, restoring the WT copy of *umpA* in colony 11 also restored the tolerance to the levels of Δ*ingB*. Taken together, these data identify a new antifungal resistance factor, *umpA,* in pathogenic molds and suggest that tolerance may lead to the emergence of azole drug resistance.

## Discussion

Studies from bacterial species have recognized antibiotic tolerance as a significant driver of treatment failures, particularly in chronic infections (40). Antibiotic tolerance also serves as a stepping stone to the development of acquired resistance (15, 16). Despite recent studies focused on identifying the genetic drivers of azole tolerance (41, 42), a significant gap remains in our understanding of how *A. fumigatus* survives or even grows under high azole concentrations and how this may impact treatment outcomes. Addressing these clinically relevant questions is expected to aid in the development of novel antifungal strategies to mitigate persistent infections in chronic aspergillosis.

We identified IngB as a novel regulator of azole tolerance whose loss may quickly lead to true azole resistance under azole selective pressure. The azole tolerance in Δ*ingB* is not dependent on morphology or developmental stage. Importantly, the *in vitro* phenotype translated *in vivo* to a murine model of invasive pulmonary aspergillosis. What is intriguing is the increase in Δ*ingB* fungal burden upon azole treatment compared to the control strains. These data raise the possibility that azole tolerant strains in human infections may be more difficult to eradicate and this warrants further study. Support for this hypothesis comes from the emergence of a putative loss of function *ingB* in *A. fumigatus* isolates longitudinally isolated from a patient with chronic granulomatous disease. Further studies are needed to determine the role of these tolerance promoting mutations in persistence in long-term chronic infections.

An outstanding question of course is how loss of *ingB* promotes azole tolerance. Loss of *ingB* had a direct effect on transcript levels of genes involved in the ergosterol pathway, iron starvation, and secondary metabolism (**Fig. 3A**). Consequently, genes critical for normal metabolic flux through pathways feeding into ergosterol production were perturbed in the absence of *ingB.* Previously, in *A. fumigatus,* a critical metabolic link between the ergosterol pathway and the siderophore biosynthesis pathway was investigated (37). Under iron starvation, mevalonate is diverted towards siderophore biosynthesis, which results in reduced metabolites for the ergosterol pathway. Based on this and the transcriptome data in the *ingB* mutant strain, we hypothesized that loss of *ingB* results in metabolic rewiring that reduces flux through the ergosterol biosynthesis pathway. Consistent with our hypothesis, Δ*ingB* showed tolerance to other ergosterol pathway inhibitors and an increase in TAFC production. Surprisingly, the expression of transcription factors HapX, SreA, or SrbA, known regulators of iron starvation and the ergosterol pathway (43–45), were not significantly different in *ΔingB*. It is possible that the chromatin structure in Δ*ingB* is altered due to reduced acetylation which affects their basal transcription levels.

While a reduction in ergosterol biosynthesis flux is the most parsimonious hypothesis to explain the Δ*ingB* tolerance, our data suggest possible alternatives. For example, *hsp90* gene expression was increased by approximately 2-fold (**Supplementary Table 1**). Although in *A. fumigatus* heat shock proteins primarily mediate echinocandin paradoxical effect and susceptibility, their involvement in azole tolerance here cannot be completely ruled out (30). In *Candida* spp. Hsp90 mediates azole tolerance and resistance (46, 47). We also observed a modest increase in the expression of *mdr1* and *atrF* transporters, which are typically overexpressed in azole resistant strains (32, 33). These data suggest there could be multiple contributing factors to the observed tolerance in Δ*ingB* to azoles.

The phenomena of azole tolerance and resistance have been studied separately; the relationship between them is a significant knowledge gap in *A. fumigatus*. Our data show that a single exposure of a tolerant strain, but not a sensitive strain, to a high azole concentration is sufficient to develop azole resistance in *A. fumigatus* (**Fig. 4**). Hence, tolerance is not merely a survival mechanism, but it also plays an active role in shaping antifungal resistance. A limitation of our study here is that we can only state that IngB mediated azole tolerance promotes resistance. It will be interesting to test non-IngB dependent tolerant strains to see if azole resistance also arises in those ill-defined genotypes. Specific metabolic perturbations that occur in the absence of *ingB* may lead to the promotion of azole resistance. These future studies will inform not only mechanisms of tolerance, but a further understanding of the biochemical pathways mediating azole resistance.

In *Candida albicans*, mechanisms of drug tolerance and resistance are parallel mechanisms and occur at different drug concentrations, and a transition from tolerance to resistance does not happen (14). Azoles, unlike their fungicidal activity against *A. fumigatus*, are fungistatic against *C. albicans.* It is possible that fungistatic drugs do not generate the same selection pressure as fungicidal drugs at high drug concentrations. As echinocandins such as micafungin are fungistatic against *A. fumigatus*, it would be interesting to test whether a link between tolerance and resistance exists here. Alternatively, passaging *A. fumigatus* at sub-MIC concentrations of azoles may also reveal the necessity of tolerance in evolution of resistance.

We identified a novel frameshift mutation in Afu5g10740 (*umpA*) that leads to azole resistance. UmpA is highly conserved in fungi but has not been studied to date in *A. fumigatus.* In *S. cerevisiae,* it is required for the correct maturation of the 20S proteasome for non-lysosomal protein degradation (48). Given the nature of the mutation, we hypothesized it to be a loss-of-function mutation, and indeed, both *umpA^ser55fs^*and the Δ*umpA* strains are resistant to azoles. On solid agar plates at 4x MIC, *umpA^ser55fs^* and the Δ*umpA* show a reduction in colony size by about 40%; however, Δ*ingB umpA^ser55fs^* (colony 11) shows no reduction at 4x, suggesting that epistatic interactions occur between *ingB* and *umpA* under azole stress.

Rpn4 is a transcription factor in *C. albicans* that activates proteasome genes to overcome the proteotoxicity induced by fluconazole (49). Loss of *rpn4* results in a loss of azole tolerance. Even in *A. fumigatus*, exposure to azoles causes oxidative and proteotoxic stress (50), and the damaged proteins are likely to be removed through ubiquitin-mediated degradation. This raises the question: How is Δ*umpA* or *umpA^ser55fs^*resistant to azoles? In *Saccharomyces cerevisiae,* the proteasome is also involved in maintaining genomic stability (51). Loss of *ump1* results in a hypermutator phenotype, characterized by an increased frequency of spontaneous mutations (52, 53). This is because proteasome activity is reduced in the Δ*ump1* strain, which prevents the rapid degradation of mutagenic DNA repair factors, specifically components of the Translesion Synthesis (TLS) pathway such as polymerase Pol 𝜁. In *A. fumigatus*, hypermutator strains resulting from impaired DNA mismatch repair increase the frequency of azole resistance (39, 54). Hence, it is possible that *umpA^ser55fs^* mutation and Δ*umpA* could also result in a hypermutator phenotype that could further drive antifungal resistance. Slower degradation of the drug target Cyp51A/B or hyperactivation of a stress response pathway in the presence of azoles could also drive resistance in Δ*umpA*.

In conclusion, we identified IngB as a novel regulator of antifungal azole tolerance and its loss leads to development of azole resistance. The null mutant is characterized by increased tolerance, accompanied by reduced expression of genes in the ergosterol pathway and increased expression of genes involved in drug efflux and the iron starvation response (**Fig. 4F**). Our study further showed that tolerance accelerates the selection of acquired resistance (**Fig. 4F**). Our future work aims to determine how UmpA, a putative 20S proteosome maturation factor, drives azole resistance. Thus, our study not only characterized a novel regulator of azole tolerance but also addressed the missing link between tolerance and resistance. Future antifungal therapies designed to reduce drug tolerance could be an effective strategy to reduce the emergence of antifungal resistance.

## Materials and methods

### Strains and culture

The reference strain AF293 (55) was used as the background strain to generate mutants. The list of strains used in the study is provided in the supplementary data. The cryostocks were maintained in 25% glycerol at −80℃. Three days prior to the experiment, the strains were streaked on Glucose Minimal Media (GMM) agar with trace elements and salt solution (56), and the plates were incubated at 37 °C, 5% CO_2_. The conidia were collected in 0.01% tween-80 water and filtered through autoclaved Miracloth. All experiments in the study were performed in liquid or solid GMM.

### Generation of Aspergillus fumigatus strains

Both *ingB* and *umpA* null mutants were generated using CRISPR-Cas9 technology as previously described (57). Two gRNA targeting sequences, one upstream and one downstream of the gene, were designed. For transformation, AF293 protoplasts (58) were transformed with a preformed Cas9-RNP complex (containing Cas9 and gRNA) and 2 μg of a repair construct. The repair construct consisted of a hygromycin expression cassette with 35-bp overhang sequences adjacent to the Cas9-targeted sites. The transformants were selected on glucose minimal media (GMM) with 1.2 M sorbitol (SMM) and hygromycin (175 μg/mL). The mutants were confirmed by junction PCRs and by checking for the absence of the open reading frame in the mutant. RT-qPCR was also used to confirm the expression in the null mutants and the reconstituted strains described below.

For the generation of reconstituted strains, *the ingB* or *umpA* gene locus (sequences obtained from FungiDB) was amplified from the AF293 genome. The resulting product was integrated into a plasmid containing pyrathimaine (*ptrA*) selection marker using HiFi (NEB), such that the selection marker lies downstream to the gene locus. For the generation of the allele swap strain, the *umpA^Ser55fs^* gene locus was amplified from colony 11, and the plasmid was generated similarly. The constructs were amplified from the plasmids using PCR and were integrated at the *aft4* safe haven site (59). The integration at the locus was confirmed using junction PCR and checking for the presence of ORF. The list of primers and gRNAs used in the study are described in **Supplementary Table 4**.

### Azole susceptibility experiments

For the solid plate assay, 1000 conidia were spotted in the middle of the plate containing various concentrations of azoles. After 72 hrs, the radial growth was measured, and the plate pictures were taken.

Minimum Inhibitory Concentration(s) were determined in accordance with the guidelines established in the CLSI M38 document (60), with the exception of using 5% CO_2_ to mimic host conditions.

To test biofilm susceptibility, 10^5^ conidia/ml were cultured in a 6-well plate for 16 hours and then exposed to azoles for 4.5 hours. The drug was removed, and the wells were washed with media, and were grown for another 16 hours. The biofilms were scraped, collected, washed, lyophilized, and weighed before and after treatments.

### Microscopy

10^5^ conidia/ml of each strain were cultured in ibidi 8-well glass bottom slides for 16 hours. The biofilms were stained with 25 µg/ml Calcofluor white for 15 minutes. The images were taken at 20X using an Andor W1Spinning Disk Confocal microscope set up on a Nikon Eclipse Ti inverted microscope and were processed using Fiji.

### Murine experiments

The animal protocol was approved by the Institutional Animal Care and Use Committee (IACUC; federal-wide assurance number A3259-01) at Dartmouth College. The experiments were conducted with strict adherence to the recommendations outlined in *the Guide for the Care and Use of Laboratory Animals* (61).

20-24g outbred female CD-1 mice (Charles River Laboratory, Raleigh, NC, USA) were housed in autoclaved cages (3-4 per cage) with HEPA-filtered air, food, and autoclaved water available *ad libitum*. The mice were given 50% grapefruit juice (62–64) 5 days before inoculation, instead of drinking water in their bottles. For immunosuppression, the mice were intraperitoneally injected with 150 mg/kg cyclophosphamide on two days before and three days after inoculation. The mice were also subcutaneously administered with 40 mg/kg Kenalog-10 (triamcinolone acetonide; Bristol-Myer Squibb, Princeton, NJ) a day before inoculation. On the day of inoculation, conidia were collected, counted, and resuspended in PBS, and 10^6^ conidia in 40 µl sterile PBS were administered intranasally. The mock group received sterile PBS. After 16 hours, the groups receiving voriconazole were given 40 mg/kg in PBS by oral gavage and once daily thereafter. At the end of the experiment, the mice were sacrificed, and their lungs were harvested, flash-frozen, and lyophilized. The relative fungal burden was assessed through quantitative PCR (qPCR) quantitation of *A. fumigatus* 18S rDNA, as previously described (65).

### RNA seq transcriptome analysis

For extracting RNA, 10^5^ conidia per ml were inoculated in petri dishes. After 16 hours, the biomass was collected in TRI reagent (Invitrogen) and bead-beaten for 1 minute. 0.2 mL of chloroform per mL of TRI reagent was added to each tube. After a brief incubation, the tubes were centrifuged for 15 minutes at top speed. The aqueous phase was collected and processed according to the manufacturer’s instructions (RNeasy, Qiagen). After DNase treatment, the quality of RNA was determined by gel electrophoresis, nanodrop, and Qubit. The final samples were submitted to the genomics core at Dartmouth College, which performed fragment analysis and library preparation for PolyA RNA-seq.

The resulting raw reads quality was checked using Initial QC. The raw reads were trimmed for adapters using CutAdapt (66). The reads (∼ 10 million per sample) were then aligned to the AF293 genome obtained from FungiDB, v68 (67) using the STAR package (68). On average, 90% of the reads aligned to the genome. The counts for each gene were compiled using HTSeq software (69). The custom R scripts used for exploratory analysis, differential expression, and figure generation are derived from the pipeline previously described (70). Briefly, the counts file and metadata were used for the exploratory analysis using the DESeq2 package (71). A Variance Stabilizing Transformation (VST) was applied (72), which normalizes the count data and transforms it onto a log2 scale, making the variance largely independent of the mean and thus suitable for PCA plots. The data were then normalized using the Trimmed Mean of M-values (TMM) method (edgeR) (73), and edgeR package (74) and limma-voom pipeline (75) were used to identify statistically differentially expressed genes between the mutant and the WT. RNA seq datasets are available in the Gene Expression Omnibus under the accession number GSE314107.

### Siderophore quantification

5 x 10^7^ conidia were inoculated in baffled flasks containing 100 ml GMM (no iron in trace elements). The flasks were incubated at 37℃, 5% CO2, and 200 rpm. After 48 hrs, the biomass was collected, lyophilized, and weighed. The supernatants were filtered through a 0.45-micron filter, and the pH was measured to ensure it was close to 6.5. To 95 μL of supernatant, 5 μL containing 30mM FeSO_4_. 7H_2_O in 5mM HCl solution were added. To control, only 5 mM HCl solution was added. After 30 minutes, the absorbance was measured at 440 nm (where TACFC production is detected maximally). The media only controls (with and without iron) were used to ensure the change of color to orange was observed in fungal samples, and not due to the background. The absorbance for each sample with 30mM iron was subtracted from the sample with solvent only. The result was divided by their respective biomass.

### Whole Genome Sequencing

All strains used for the WGS were prepared by culturing 10^5^ conidia per ml in petri dishes. After 24 hrs, the biomass was collected, frozen, and lyophilized. The resulting biomass was subjected to a bead beater for 1 minute, and DNA was extracted using previously published protocols. The samples were submitted to Seqcoast (Portsmouth, NH) for short-read Illumina sequencing and variant calling. Samples were prepared for whole genome sequencing using the Illumina DNA Prep tagmentation kit (#20060059) with Illumina Unique Dual Indexes. Sequencing was performed on the Illumina NextSeq 2000 platform using a 300-cycle flow cell kit to produce 2x150 bp paired reads. 1-2% PhiX control was spiked into the run to support optimal base calling. Read demultiplexing, read trimming, and running analytics were performed using DRAGEN v4.2.7, an on-board analysis software on the Illumina NextSeq 2000. The whole genome sequencing data have been submitted in the NCBI repositories. The raw sequencing data is available via Sequence Read Archive (SRA) under the BioProject ID - PRJNA1379593. Detailed sample information including BioSample accession numbers and SRA run accession numbers is included in **Supplementary Table 5**.

### Mutation analysis

To identify mutations in strains the Illumina DNA sequence was aligned to the genome with bwa-mem2 v2.2.1 (76) and converted to a sorted BAM format alignment file with samtools v1.19.2 (77) and optical duplicates marked with MarkDuplicates in the Picard Toolkit (“Picard Toolkit.” 2019. Broad Institute, GitHub Repository. https://broadinstitute.github.io/picard/; Broad Institute). Variants were identified using GATK HaplotypeCaller and GenotypeGVCFs v4.6.0.0 (78). To identify impact of variants snpEff v4.3t (79) was run with the AF293 genome and annotation downloaded from FungiDB v68 (67). The resulting VCF file was processed with custom python script to generate table of variants to examine as a spreadsheet. Scripts for processing data and VCF variant call are archived in github repository https://github.com/stajichlab/CramerLab_Afumigatus_AF293_VOR and archived in Zenodo at doi: 10.5281/zenodo.17981730.

### Statistics

Unless otherwise stated all statistics and graphs are generated using Graphpad Prism (v10.6.1) from three biologically independent experiments (unless otherwise stated) with multiple technical replicates in each experiment.

## Data availability

All other data are included in the main article or in the *SI* and *Dataset S01*.

## Acknowledgements

This work was funded by NIH/NIAID grants R01AI130128 (R.A.C.) and R01AI146121 (R.A.C.). It was further supported by the Cystic Fibrosis Foundation Research Development Program (STANTO19R0) and NIH/NIDDK P30-DK117469 (Dartmouth Cystic Fibrosis Research Centre). S.V. is supported by a postdoctoral fellowship from Cystic Fibrosis Foundation (VELLAN22F0). RNA sequencing was carried out in the Genomics and Molecular Biology Shared Resource (RRID:SCR_021293) at Dartmouth which is supported by NCI Cancer Center Support Grant 5P30CA023108 and NIH S10 (1S10OD030242) awards. Authors would also like to thank Dr. Fred W. Kolling IV, Dr. Elizabeth Ann Sergison, Dr. Owen M. Wilkins, and Dr. Charles T.S. Puerner for assistance in RNA sequencing analysis. We thank Dr. Daniel Schultz for feedback on the manuscript. Authors would also like to thank Ann Lavanway in the Dartmouth Microscopy core for assistance in confocal microscopy. We thank the staff of the Center for Comparative Medicine and Research (CCMR) for providing animal husbandry and facility support for the duration of this study.

## Author Contributions

S.V and R.A.C. designed research; S.V., N.D., C.L., J.E.S.., performed research; S.V., J.E.S., and R.A.C contributed new reagents/analytical tools; S.V., N.D., C.L., J.E.S.., and R.A.C. analyzed data; and S.V, J.E.S., R.A.C. wrote the manuscript.

